# Decision analysis shows scientific and economic value of community-based monitoring in Madagascar

**DOI:** 10.1101/2024.10.06.616891

**Authors:** Lydia G. Soifer, David H. Klinges, Lalatiana Randriamiharisoa, Brett R. Scheffers

## Abstract

Community-based monitoring (CBM) could enable long-term biodiversity monitoring in remote areas and benefit local communities, but its benefits have rarely been quantified. We use a multi-criteria decision analysis framework to systematically examine the scientific and socioeconomic values and financial costs associated with biodiversity monitoring for vertebrates by scientists and local community members in six protected areas (PAs) in Madagascar, encompassing diverse ecosystems spanning tropical rainforests to spiny deserts. We compare the number of species observed during scientist and community surveys, identify the ‘ideal’ number of scientist and community surveys that would be required to maximize the scientific and socioeconomic values of monitoring efforts while minimizing financial cost, and compare monitoring plans across several conservation philosophies representing “ecocentric” and “people-centered” perspectives. Scientists generally observed more species than community members. However, including a greater proportion of surveys conducted by community members improves wildlife monitoring across PAs, taxonomic groups, and diverse conservation philosophies due to the lower financial cost of travel and compensation relative to monitoring exclusively conducted by scientists. While the valuation schemes we use are simplistic representations of the complex costs and values associated with CBM, this study indicates the benefits of community monitoring regardless of the conservation philosophy used to anchor valuation and decision-making. Increasing integration of CBM into existing conservation management could therefore offer a financially viable method to consistently monitor biodiversity and benefit local communities in the face of limited funding and global challenges.

## 1. Introduction

Ecological monitoring is essential to understand the impacts of anthropogenic and natural disturbances on biodiversity and ecosystem function and to respond to these global challenges with adaptive conservation management (Holl, 2017; Niemelä, 2000; Pereira & Davidcooper, 2006). At local scales, policy makers and protected area (PA) managers depend on data from long-term monitoring efforts to forecast population trends, identify priority areas for conservation, and inform conservation schemes (Danielsen et al., 2021; Danielsen, Jensen, et al., 2014; Lindenmayer et al., 2012; Wyborn & Evans, 2021). However, biodiversity monitoring is often overlooked during the planning phase of conservation projects, given inadequate funding and its dependence on external scientists (Danielsen et al., 2021; Lindenmayer et al., 2012). These issues often result in short-term or infrequent monitoring efforts that are unlikely to fulfill the data needs required for effective conservation decision-making (Danielsen et al., 2003). Community-based monitoring (CBM), including integration of local communities into design and implementation of monitoring plans, could help overcome these monitoring challenges by providing a financially feasible and sustainable solution (Danielsen et al., 2003, 2007; Holck, 2008).

Community-based monitoring can help inform conservation management and generate positive outcomes for conservation and communities (Danielsen et al., 2007, 2021; Dolch et al., 2015; Moller et al., 2004). For example, within communities, CBM may provide income, improve local organizational structure, and enhance community cohesion and pride of local natural resources, which together can create internal social and political pressures to reduce rates of natural resource extraction (Andrianandrasana et al., 2005; Danielsen et al., 2021; Dolch et al., 2015; Fernandez-Gimenez et al., 2008; Fry, 2011; Humber, Godley, et al., 2017; Razanatsoa et al., 2021). The socioeconomic empowerment communities gain from involvement in monitoring efforts may provide the foundation for monitoring programs to be continued by local communities once initial funding schemes have finished (Kainer et al., 2009).

While CBM is slowly being more widely integrated into conservation practice due to the range of benefits it may provide, its use for informing conservation management has been met with resistance (Carvalho et al., 2009). Reluctance by decision-makers to use community-collected data generally stems from concerns of data quality or that community members may report biased results when local interests differ from agency goals (Cretois et al., 2020; Danielsen et al., 2021; Danielsen, Jensen, et al., 2014; Nielsen & Lund, 2012). Despite these concerns, studies reporting the quantitative comparability of monitoring efforts conducted by communities and external scientists are increasing in frequency and occur across diverse geographic regions (e.g., Danielsen, Jensen, et al., 2014; Holck, 2008; Humber, Godley, et al., 2017). For agencies to adopt community-based biodiversity monitoring, the financial viability and scientific and socioeconomic values of CBM to conservation and communities must be quantified.

Multi-criteria decision analysis (MCDA) provides a quantitative framework to systematically evaluate the overall values obtained from different conservation plans based on conservation philosophies held by stakeholders (Blattert et al., 2017; Gregory et al., 2012; Uhde et al., 2015). Stakeholder philosophies may range from traditional ‘ecocentric’ conservation to new ‘people-centered’ conservation (Kareiva & Marvier, 2012; Sandbrook et al., 2019). In 2014, Georgina Mace described the traditional ecocentric conservation philosophy, which dominated viewpoints from the 1960s-1990s, as “Nature for itself” or “Nature despite people”, emphasizing the protection of nature for its own sake (Mace, 2014). Its framing is focused on well-established metrics, such as the number of species listed in the IUCN red list or the coverage of protected areas. The ecocentric philosophy thus corresponds to monitoring plans that are able to most accurately and exhaustively record species’ occurrences. In contrast, Mace (2014) describes the new ‘people-centered’ philosophy, which emerged in the early 2000s as “Nature for people” or “People and nature”. The people-centered conservation philosophy emphasizes the importance of conservation for humans, economic valuations, and socioecological systems, and would therefore value engaging local communities in biodiversity monitoring (Sandbrook et al., 2019). Using MCDA to compare different monitoring plans defined by the number of surveys performed by scientist and community members, it is possible to identify plans that minimize financial cost while maximizing the benefit of monitoring for scientific and socioeconomic objectives. These objectives can be defined by different conservation philosophies ranging across a spectrum from a fully ecocentric philosophy that favors observing the maximum number of species with little regard for the number of community members engaged in monitoring, to a fully people-centered philosophy that favors monitoring plans that provide data sufficient for PA management and that incorporate community members at a high degree of involvement.

In Madagascar, long-term biodiversity monitoring efforts are urgently needed to assess and protect the island’s largely endemic biota, which is severely threatened by habitat loss (Ralimanana et al., 2022). Though CBM has been successful in several regions within Madagascar, it has not been widely adopted throughout the country despite its potential to benefit adaptive management in remote areas where limited access has largely prevented consistent monitoring (Andrianandrasana et al., 2005; Dolch et al., 2015; Humber, Andriamahaino, et al., 2017; Humber, Godley, et al., 2017). To determine if community monitoring programs are generalizable across Madagascar’s protected areas (PAs), we examined the contribution of community-based biodiversity monitoring to conservation efforts by first comparing the vertebrate species richness (amphibians, birds, mammals, reptiles) observed during surveys conducted by community members (CMs) versus scientists in six ecologically unique PAs (Fig. 1). We then used MCDA to 1) identify the ideal number of CM and scientist surveys that would minimize financial cost while maximizing the overall value of monitoring to conservation objectives as defined by different conservation philosophies and 2) compare the value of different monitoring plans to conservation objectives and their associated financial costs under an ecocentric conservation philosophy and a people-centered conservation philosophy. Our study not only informs strategies for CBM in Madagascar, but also provides a reproducible and generalizable framework for incorporating multiple conservation philosophies into conservation project planning that can be applied anywhere globally and for projects of varying spatial and temporal scales and budgets.

**Fig. 1.**
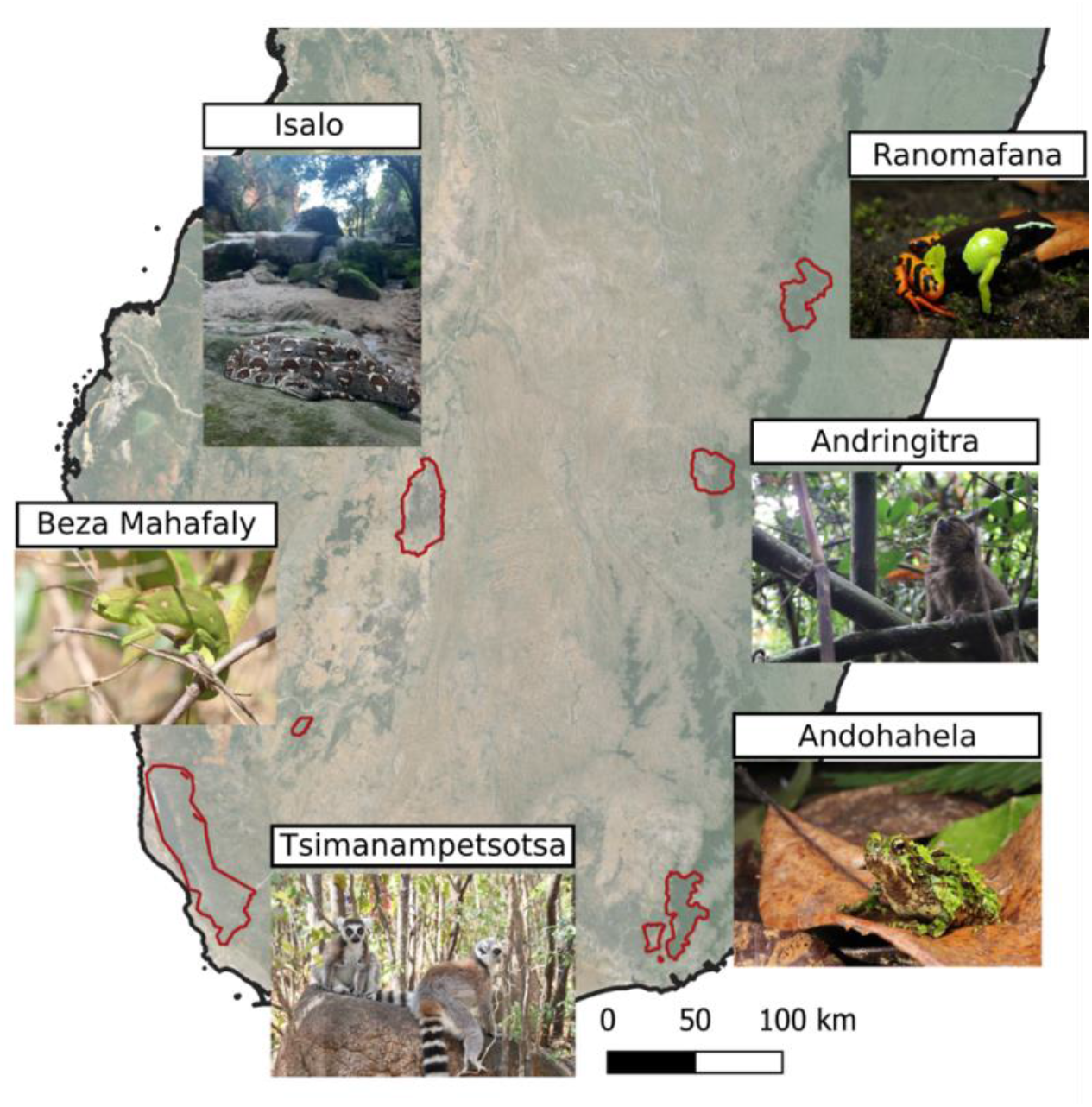
Map of southern Madagascar with protected areas used in the monitoring study. Images show a vertebrate species representative of each protected area, which ranged from dry and spiny forest (Tsimanampetsotsa, Beza Mahafaly) to transition forest (Andohahela, Isalo) to wet forest (Andringitra, Ranomafana).

## 2. Methods

### 2.1. Study site

The study took place in six protected areas (PAs) in Madagascar spanning five of the seven ecoregions in the country (Olson et al., 2004). PAs included Ranomafana National Park, Andringitra National Park, Andohahela National Park, Beza Mahfaly Special Reserve, Isalo National Park, and Tsimanampetsotsa National Park (Fig. 1).

### 2.2. Knowledge exchange workshops between scientists and community members

For each of the six PAs, we collaborated with local community members (CMs) to co-develop monitoring methods for all vertebrate taxa. We created a photographic guidebook for mammals, birds, reptiles, and amphibians for each PA based on community knowledge of the local fauna. The local and scientific names of each species were accompanied by a photograph of the species (or multiple photographs for dimorphic species), classified by researchers who worked in the PA or other Madagascar National Park staff. Such staff trained CMs in standardized data collection methods, emphasizing the importance of standardized observations and teaching them to use the guidebook, GPS devices, and data sheets. Scientists were graduate students at the University of Antananarivo who specialized in the ecology of Malagasy mammals, birds, reptiles, or amphibians. To ensure CMs were proficient at identifying species, all CMs undertook an initial training period during which scientists and CMs simultaneously conducted surveys independently on the same transect. CM and scientist surveys were then compared, and additional side-by-side surveys were conducted until the data collected by both groups were 50% similar. Training was completed over three to five sessions in three to five days depending on the number of species in the PA. Training is further described in Price et al. (2023).

### 2.3. Data Collection

Surveys were conducted in 2019 and 2020 in the six PAs (**Error! Reference source not found**.). In each PA, a minimum of six 1,250 m transects were established in areas that maximized the variability of habitats being monitored. Additionally, two transects were established near bodies of water to improve detection of amphibians. All scientist and CM surveys occurred in the same months and during the daytime to eliminate potential bias due to seasonality or circadian cycles. In total, we had 148 surveyors comprising 19 scientists and 129 CMs who conducted 1763 surveys across the six PAs; 1,281 surveys were conducted by community members and 482 surveys were conducted by scientists (Table S1). These survey totals exclude training surveys where CM and community members conducted surveys side-by-side.

### 2.4. Comparing scientist and community surveys

To compare rarefied species richness observed by scientist and CM surveys for each taxonomic group in each PA, we calculated sample-based species accumulation curves for CMs and scientists using 100 permutations. For each taxonomic group (mammals, birds, reptiles, or amphibians) in each PA, we then compared the number of species seen in surveys conducted by scientists or CMs, rarefying survey effort to the lower maximum number of surveys conducted by scientists or CMs. For example, if scientists conducted six surveys and CMs conducted twelve surveys, we compared the average number of species seen in six surveys conducted by scientists versus six surveys conducted by CMs.

### 2.5. MCDA framework

We used the MCDA framework to compare the value of monitoring efforts to different conservation philosophies and financial costs associated with different monitoring plans (Fig. 2, 3). We defined monitoring plans based on the number of scientist and CM surveys (Table 1). By comparing many monitoring plans, we first identified the optimal number of CM and scientist surveys for each taxonomic group within each PA that would maximize the value of monitoring to different conservation philosophies while minimizing cost. We then more closely examined the value and financial costs of four monitoring plans for ecocentric and people-centered conservation philosophies.

**Table 1.** Terminology used in the MCDA framework.

| **Term** | **Definition** |
| --- | --- |
| Attribute | A component of monitoring efforts that impacts the outcome (e.g. # of species seen, local community engagement) |
| Attribute weight | A number between 0 and 1 that represents the relative importance of observing biodiversity vs. including CMs in monitoring efforts. The two weight parameters in the mathematical definition of the conservation perspective always sum to one. |
| Conservation philosophy | A perspective held by a stakeholder regarding the value and relative importance of observing species richness and community engagement to monitoring outcomes. Mathematically defined by a set of value functions and weight attributes. |
| Ecocentric philosophy | A conservation philosophy that emphasizes the protection of nature for its own sake |
| Linear additive model | Quantifies the overall value of a monitoring plan |
| Monitoring plan | The number of scientist and CM surveys |
| People-centered philosophy | A conservation philosophy that emphasizes the importance of conservation for humans, economic valuations, and socioecological systems |
| Value function | A function (e.g. $f(x) = x$) used to describe the relationship between an attribute measure and the value of the attribute to the overall value of monitoring efforts. |

**Fig. 2.**
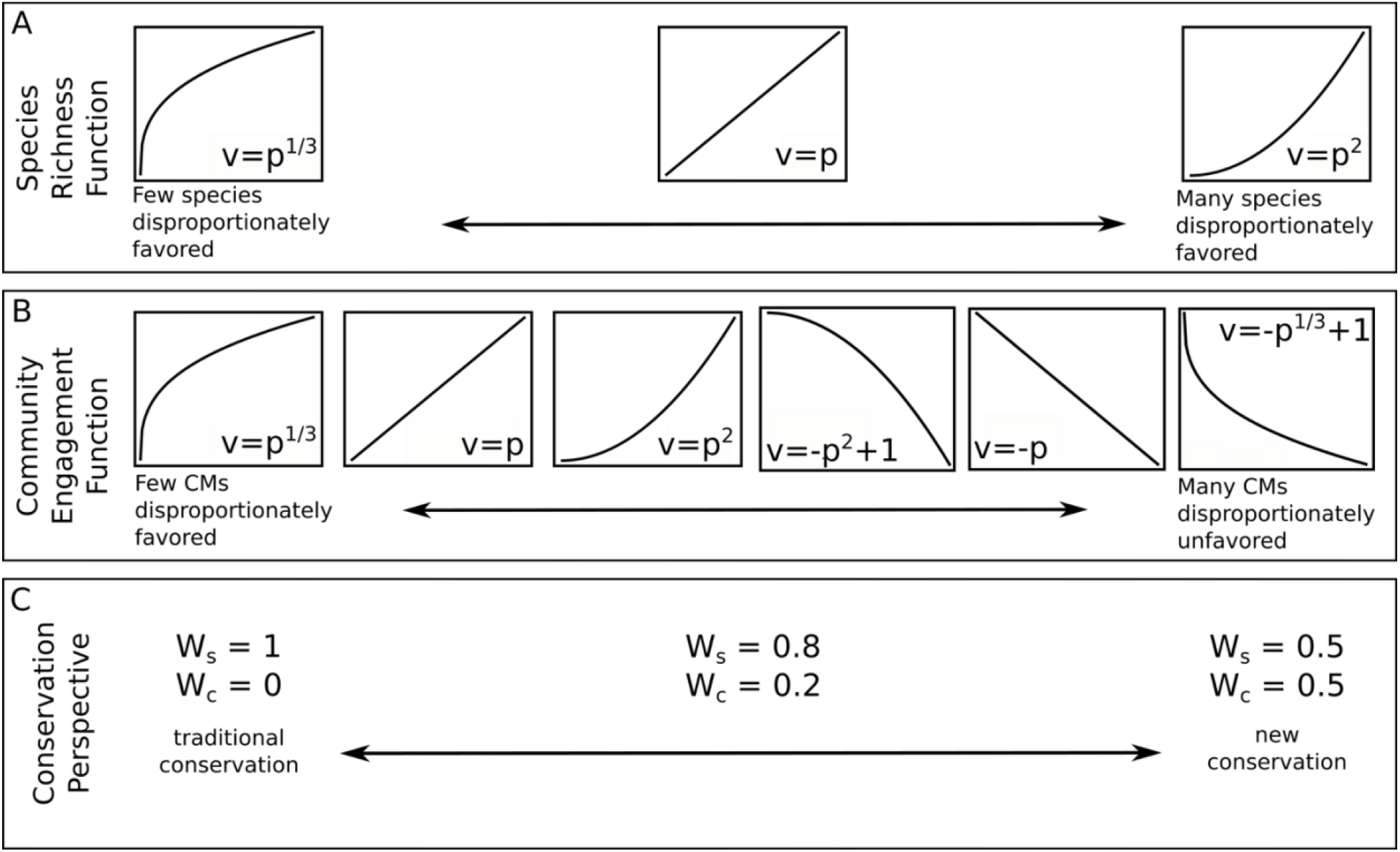
Conservation philosophies are defined by three axes. A) The species richness value function represents the benefit of observing few or many species. B) The community engagement value function represents whether many or few CM surveys benefit or impair the outcomes of monitoring efforts. C) The pair of attribute weights represent the relative importance of observing species (W_s_) versus community engagement (W_c_). Both the x and y axes of the value functions range from zero to one. The x-axis represents the standardized attribute measure (i.e., proportion of observed species (A) or proportion of community member surveys (B)). The y-axis represents the attribute value. The shape of the curve represents the perceived relationship between attribute value and standardized attribute measure.

**Fig. 3.**
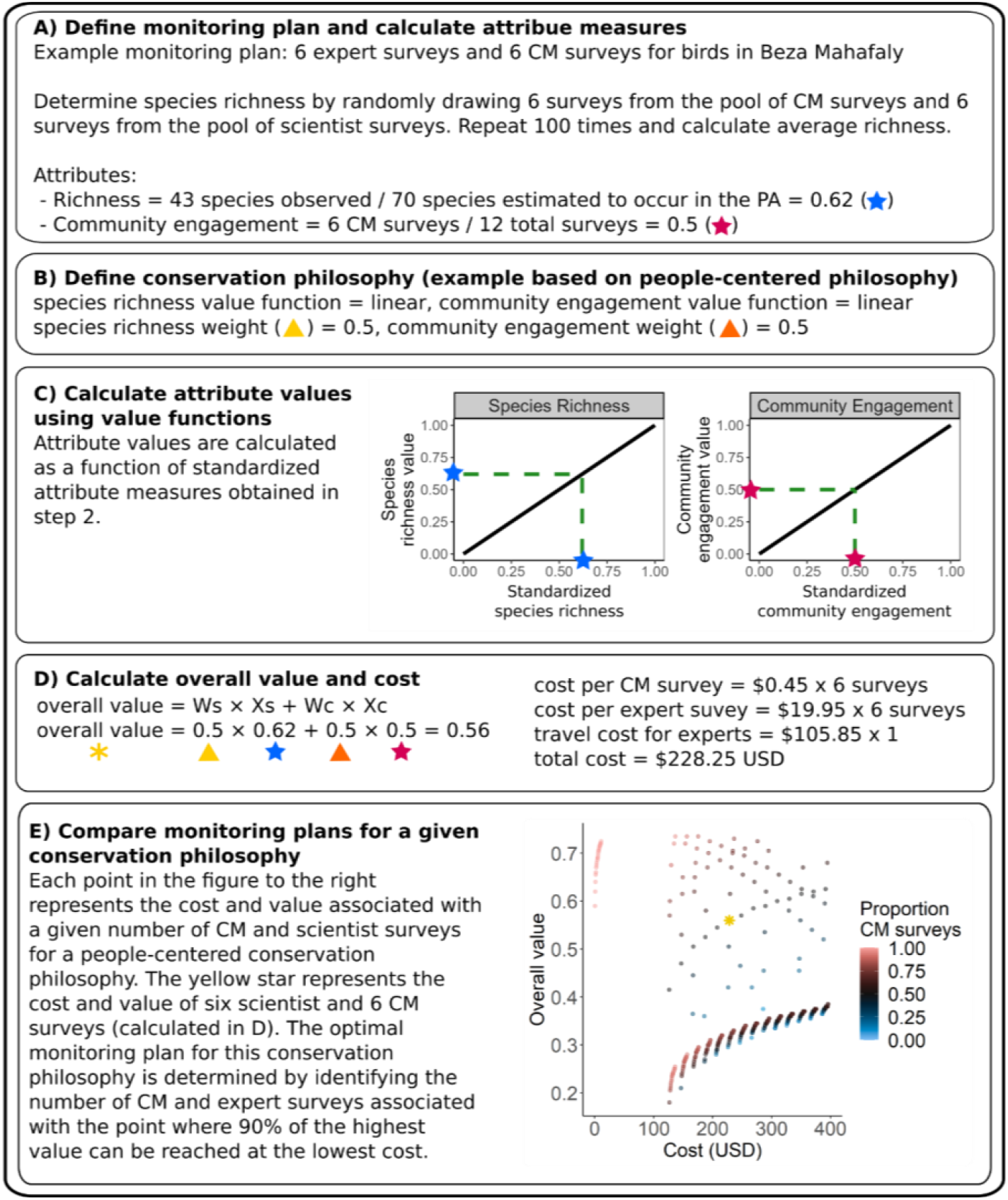
Example of methods used to quantify the overall value of a monitoring plan. Here, we show calculations for a people-centered conservation philosophy that assigns equal attribute weights and linear value functions for species richness and community engagement (as shown in Fig. 2: species richness function: v = p; community engagement function: v = p; conservation perspective: W_s_ = 0.5, W_c_ = 0.5). This example focuses on bird surveys in Beza Mahafaly with a monitoring plan defined by 6 scientist and 6 CM surveys. In step E, monitoring plans are compared that use pairwise combinations of 1 to 24 CM and scientist surveys. These steps can be repeated for any conservation philosophy and monitoring plan.

#### 2.5.1 Cost of monitoring plans

For each PA, we determined the financial cost of scientist and CM surveys based on the Madagascar National Park budget for CM and scientist surveyor salaries and transportation costs (transportation costs are only relevant to scientists as all CMs lived within walking distance of PAs). We derived a per survey cost for each PA by dividing the budget for surveyor salaries by the median number of CM and scientist surveys conducted in all PAs during 2019 and 2020 (**Error! Reference source not found**.).

#### 2.5.2. Calculating the value of monitoring plans

The value of monitoring is defined using a linear additive model (Gregory et al., 2012), which is calculated as the sum of weighted attribute measures that have been standardized and adjusted to reflect desirability of an outcome using value functions; *overall value = W*_*1*_*X*_*1*_ *+ W*_*2*_*X*_*2*_ *+ W*_*n*_*X*_*n*_, where *X*_*i*_ represents the attribute value and *W*_*i*_ represents the attribute weight (Table 1). The attribute value is calculated by standardizing the attribute measure using a mathematical function (i.e., the value function; Fig. 2A, B).

We define two attributes that influence monitoring outcomes: species richness and community engagement. For a given monitoring plan (i.e., number of scientist and CM surveys), species richness was measured as the average number of species seen across 100 permutations of randomly drawn surveys, divided by the total richness in a PA as derived from accumulation curves (i.e. the proportion of total species observed). Community engagement was measured as the proportion of surveys conducted by CMs (as opposed to scientists) for a given monitoring plan (Fig. 3A). We estimated the total number of species for each taxonomic group in each PA based on all scientist plus CM surveys using the ‘estimateR’ function from the ‘vegan’ R package (Oksanen et al., 2022) (Fig. S1-4).

We quantitatively define a conservation philosophy using a set of mathematical value functions and weights for each attribute (i.e., species richness and community engagement) (**Error! Reference source not found**., 3B).

Mathematical functions (e.g., linear, quadratic, etc.) are used to calculate the value of attributes as a function of the attribute measure (Fig. 3C). For example, a linear function would indicate that a 5% increase in the proportion of species observed equals a 5% increase in the value of observing species. A quadratic function, on the other hand, describes diminishing returns, such that a 5% increase in the proportion of species observed may equal a 10% increase in the value of observing species when relatively few species have been observed, but a 2% increase in the value of observing species once many species have been observed.

Attribute weights determine the relative importance (low to high) of each attribute (i.e., the relative importance of observing species versus community engagement). For example, a fully ecocentric philosophy highly values observing species and is ambivalent to community engagement, and therefore we would assign an attribute weight of 1 to the proportion of species observed and 0 to community engagement (Fig. 2C). Attribute values are then multiplied by attribute weights and summed to derive an overall value for a monitoring plan (Fig. 3D). The sum of all attribute weights always equals one, so that the maximum value of a monitoring plan is one and the minimum value of a monitoring plan is 0.

### 2.6. Identifying the optimal number of scientist and CM surveys across conservation philosophies

We determined the ideal number of CM and scientist surveys using the MCDA framework across 54 different conservation philosophies that are defined by attribute weights and mathematical functions and that vary in three ways relevant to conservation management: 1) the value placed on observing the first species, which are likely to be common, versus observing later species, which are likely to be rare (“species richness axis”); 2) whether increased community engagement is valued positively or negatively (“community engagement axis”); and 3) the relative importance of species richness versus community engagement (“conservation perspective axis”) (Fig. 2).

We represent the species richness axis using three monotonically increasing value functions: quadratic, linear, and exponential. The quadratic value function places greater value on observing common species, since the value of observing species increases rapidly with the first few species observed. The exponential function places greater value on observing rare species, since the value of observing few species is initially low and increases more rapidly as more species are seen. We represent the community engagement axis using six value functions that range from an exponential increase in value with increasing CM surveys (i.e. more community engagement favored), to an exponential decrease in value with increasing CM surveys (i.e. CMs impair the outcomes of monitoring efforts). We represent the conservation perspective axis using three sets of attribute weights: weight of species richness (W_rich_) = 1 and weight of community engagement (W_cm_) = 0 (i.e., an ecocentric conservation philosophy), W_rich_ = 0.8 and W_cm_ = 0.2 (i.e., a philosophy in between ecocentric and people-centered), and W_rich_ = 0.5 and W_cm_ = 0.5 (i.e., a people-centered philosophy). We derived the 54 conservation philosophies from all possible combinations of value functions and attribute weights across the three axes (3 × 6 × 3 = 54).

To identify the optimal number of CM and scientist surveys for a given conservation philosophy, we first calculated the overall values and financial costs associated with monitoring plans for all pairwise comparisons of CM and scientist surveys up to a total of 24 scientist and 24 CM surveys within a budget period. For example, monitoring plans included 1 CM and 1 scientist survey, 1 CM and 2 scientist surveys, …, 24 CM and 1 scientist survey, …, 24 CM and 24 scientist surveys, etc. We did not evaluate monitoring plans that included more surveys than were conducted for a given taxonomic group in a PA during the study. From these monitoring plans, we identified the monitoring plan with the “optimal overall value”: the plan with the lowest cost yet still with a value within 0.01 of the maximum value attained of any plan for that conservation philosophy. This represents the potential for stakeholders to be willing to compromise 1% of the overall value to reduce financial costs of monitoring. We then determine the number of CM and scientist surveys required to achieve the optimal overall value at the lowest financial cost.

### 2.7. Comparing monitoring plans based on ecocentric and people-centered conservation philosophies

We showcase how MCDA may be used in Madagascar to compare four monitoring plans within two of the conservation philosophies representing an ecocentric and people-centered perspective (Holmes et al., 2017; Sandbrook et al., 2019) (Table 1). The first monitoring plan was a “business as usual” plan composed of 6 scientist surveys and 0 CM surveys, as this most closely reflects current monitoring efforts in many Malagasy PAs. The other three monitoring plans represent increased monitoring effort, including 1) a scientist-only monitoring plan composed of 12 scientist surveys and 0 CM surveys; 2) a community-based monitoring plan composed of 12 CM surveys and 0 scientist surveys; and 3) a joint monitoring plan composed of 6 scientist and 6 CM surveys. In six instances for select taxa, scientists conducted fewer than six or twelve surveys due to logistical constraints (Table S1). We therefore excluded these cases from the comparison of monitoring plans. We define the ecocentric conservation philosophy using a linear function and attribute weight of one for species richness and an attribute weight of zero for community engagement weight = 0 (and therefore no community engagement value function). We define the people-centered conservation philosophy using a linear function and attribute weight of 0.5 for both species richness and community engagement.

## 3. Results

### 3.1. Richness observed during scientist vs. CM surveys

We compared rarefied richness between scientist and CM surveys (**Error! Reference source not found**.). Whether scientists or CMs observed more species varied across PAs and taxa. Overall, scientists observed more species in 13 of the 24 combinations of PAs and taxa. The difference in observed richness between scientists and CMs was generally low for mammals, moderate for reptiles and amphibians, and high for birds (Fig. 2). For example, the average difference in the number of mammal species seen by scientists and CMs was less than five species across PAs, while scientists observed an average of 23.36 more bird species in Beza Mahafaly.

### 3.2. Identifying ideal numbers of CM and scientist surveys across conservation philosophies

For each taxon in each PA, we calculated the number of surveys conducted by CMs and scientists required to achieve the optimal value to conservation objectives defined by 54 different conservation philosophies. Across PAs and taxa, most conservation philosophies required a combination of surveys conducted by CMs and scientists to attain the optimal value (Fig. 5). In 67% of the scenarios spanning PAs, taxa, and conservation philosophies, the optimal value required more surveys to be conducted by CMs than scientists. However, the relative number of CM and scientist surveys varied widely by PA, taxon, and conservation philosophy. For example, when conservation philosophies disfavor surveys conducted by CMs (i.e., community engagement is assigned a negative function), the monitoring plan required to attain the optimal value consists only of surveys conducted by scientists. Additionally, in Ranomafana, more scientist than CM surveys were required to attain the optimal value in 60.65% of the conservation philosophies (Fig. S5). In other PAs, across conservation philosophies that are either ambivalent to or value community engagement, the optimal value is attained with a combination of CM and scientist surveys composed of a higher proportion of CM surveys (Fig. 5).

**Fig. 4.**
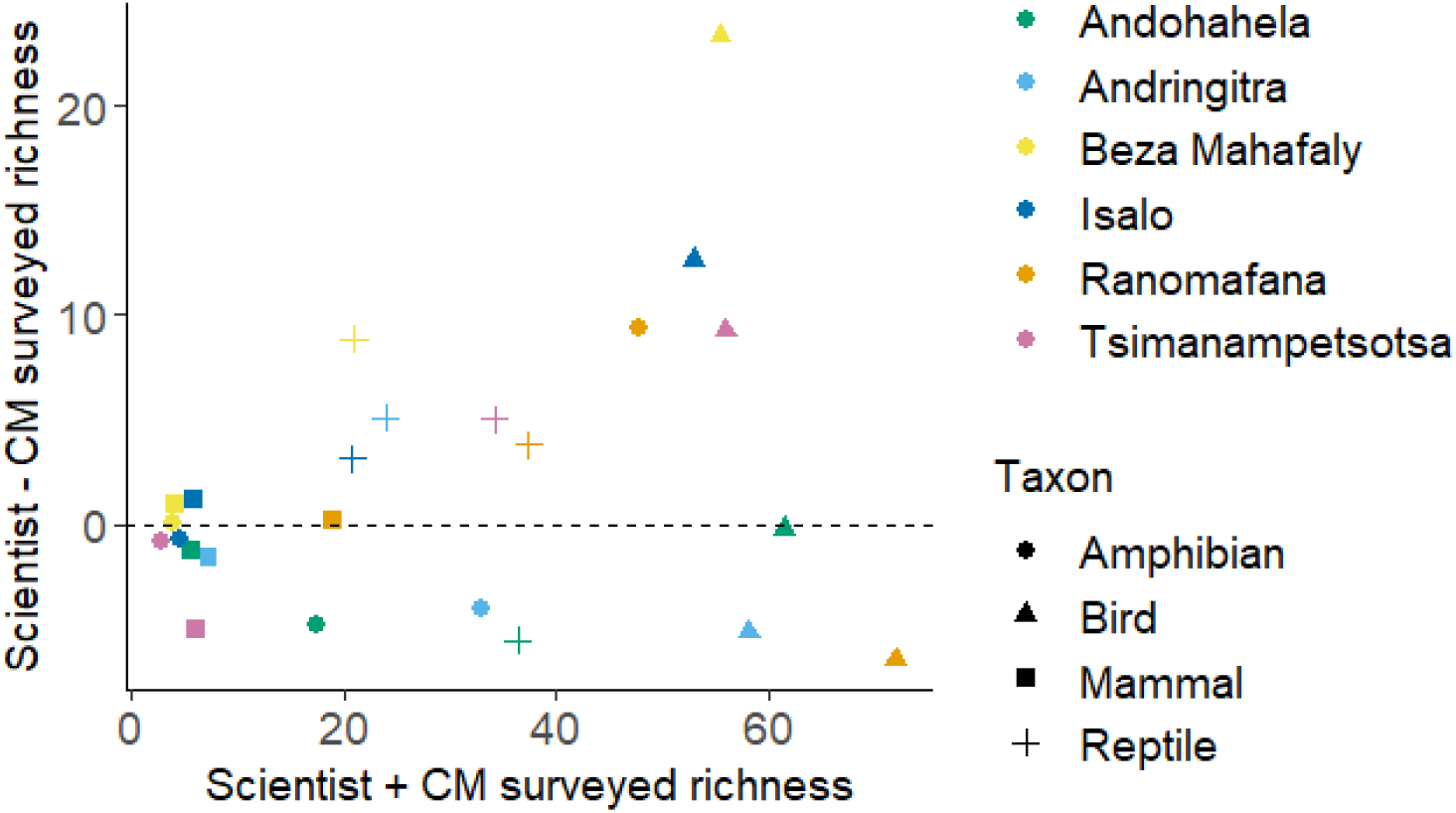
The difference between rarefied species richness from surveys conducted by scientists versus community members (y-axis) as a function of the total rarefied species richness from surveys conducted by both scientists and community members (x-axis). Rarefied richness was calculated from 100 permutations. Richness was rarefied to the lower maximum number of surveys conducted by scientists or CMs for each taxon in each PA. Higher total rarefied richness generally entailed that scientists observed more species than community members.

**Fig. 5.**
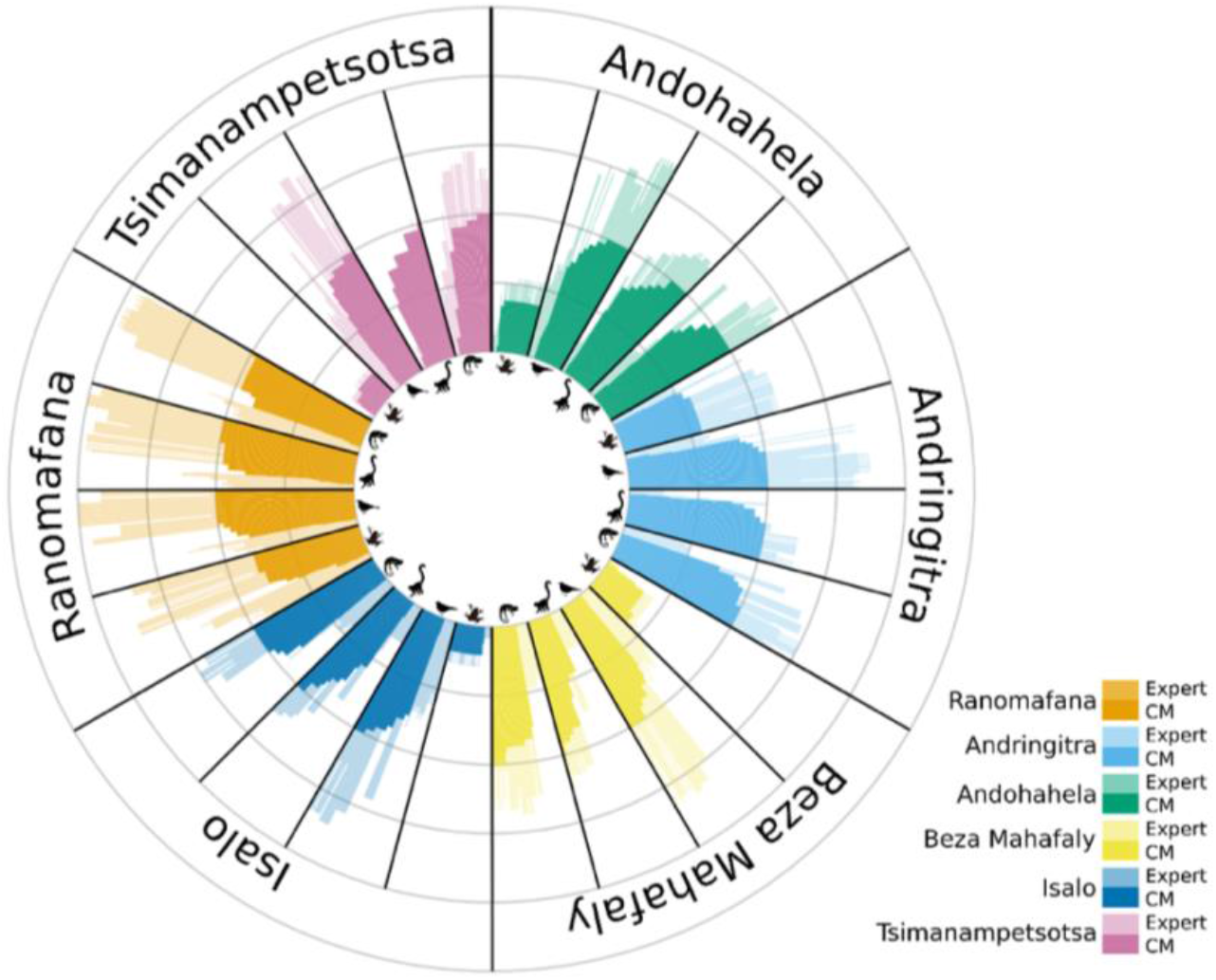
Stacked bar plot representing the total number of scientist and CM surveys required to attain the optimal overall value at the lowest cost for each PA, taxonomic group, and conservation philosophy. For each combination of PA and taxonomic group, 54 different conservation philosophies were evaluated. Within each PA, categories in the clockwise direction represent taxa: amphibians, birds, mammals, and reptiles. Within each taxon in each PA, bars represent conservation philosophies. Solid bars represent the number of CM surveys (from 0 to 24 surveys) and transparent bars represent the number of scientist surveys (from 0 to 24 surveys). Gray concentric circles indicate intervals of 12 surveys. The number of CM and scientist surveys cannot exceed the number of surveys that were conducted during the study. Across most PAs, taxa, and conservation philosophies, the combination of solid and transparent bars suggests that both CM and scientist surveys are required to attain the optimal value of monitoring.

### 3.3. Comparing monitoring plans under two conservation philosophies

We evaluated four monitoring plans that use different combinations of scientist and CM surveys based on one ecocentric and one people-centered conservation philosophy (**Error! Reference source not found**.). Within the people-centered conservation philosophy, monitoring plans using only CM surveys consistently attained the highest overall value for the conservation objectives and had the lowest financial cost across PAs and taxonomic groups. In contrast, the optimal value under an ecocentric conservation philosophy was not always attained with a CM-only monitoring plan (Fig. 6). For example, the value of monitoring efforts was 22% and 30% greater when scientists conducted twelve bird surveys in Isalo and Beza Mahafaly, respectively, than when CMs conducted twelve bird surveys in the same PAs. However, in Ranomafana, Andringitra, Andohahela, and Tsimanampetsotsa the difference between the value of monitoring efforts for birds between the scientist-only plan and the CM-only plan varied by less than 3%, 4%, 2%, and 12%, respectively (Fig. S6).

**Fig. 6.**
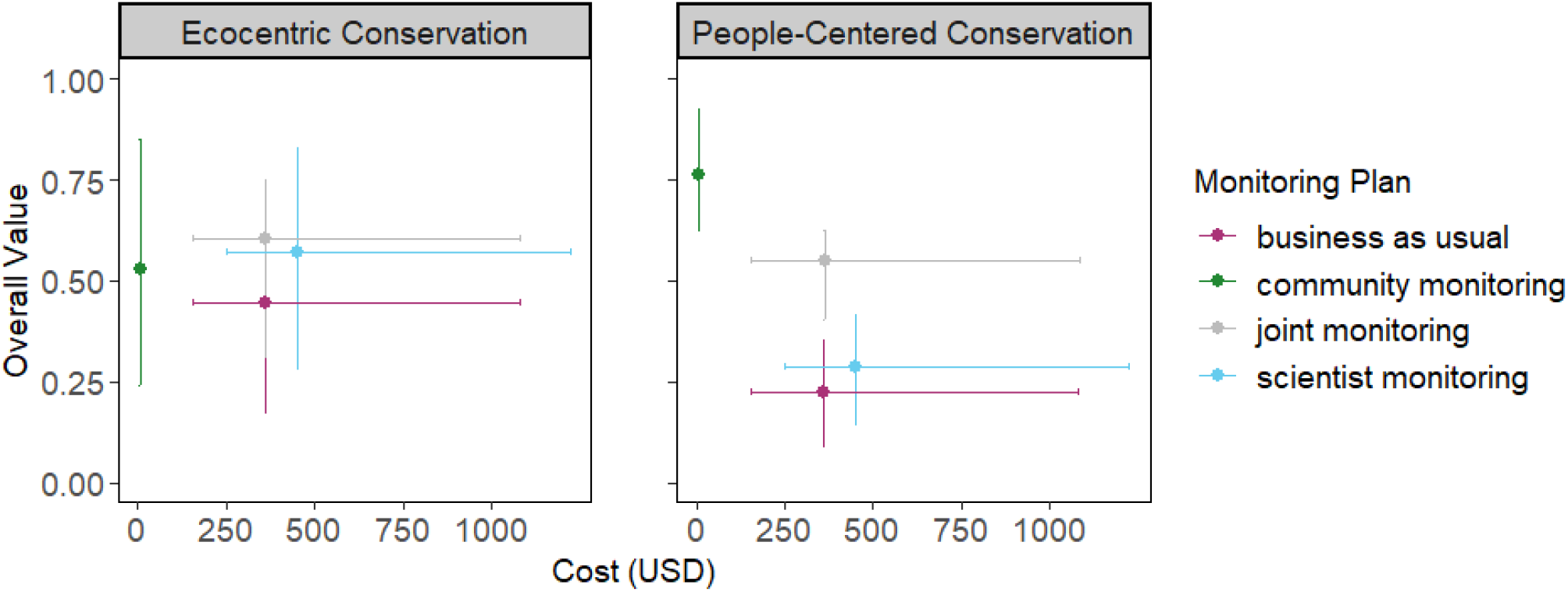
Overall costs and values associated with four monitoring plans for two different conservation philosophies. Points and error bars represent the mean and standard deviation for costs and values of all taxa and protected areas (PAs). The ecocentric conservation philosophy is defined by a species richness weight of 1, a linear species richness function, and a community engagement weight of 0. The people-centered conservation philosophy is defined by a species richness weight of 0.5, a linear species richness function, a community engagement weight of 0.5, and a linear community engagement function. The four monitoring plans include business as usual (which mostly closely matches monitoring in these PAs prior to this study: 6 scientist surveys), community engagement (6 scientist and 6 CM surveys), community monitoring (12 CM surveys), and scientist monitoring (12 scientist surveys). Overall monitoring costs and values are calculated per PA and taxon (represented by point shape – amphibians, birds, mammals, and reptiles). If the monitoring plan contains more surveys than were conducted during the study (six or twelve surveys depending on the monitoring plan), then it was excluded from the analysis.

## 4. Discussion

Defining a clear monitoring scheme during the planning phase of a conservation project is essential for successful management of biodiversity. While monitoring has often been conducted by scientifically trained experts, recent trends toward co-management have led to questioning whether monitoring efforts by local communities would produce equivalent outcomes (Danielsen et al., 2021; Gardner et al., 2018). Though comparisons of community member and scientist monitoring remain limited, case studies of communities around the globe have found that they provide relatively equivalent data (Danielsen, Jensen, et al., 2014; Holck, 2008; Price et al., 2023). Here, we show that despite variability regarding whether scientists or community members (CMs) observe more species, integrating CMs into biodiversity monitoring efforts provides a more cost-effective monitoring plan. Furthermore, we show that incorporating CMs produces outcomes with high value for conservation objectives even when based upon divergent conservation philosophies that differ in how they regard the relative importance of community engagement and observed species richness.

### 4.1. Do we need scientists for biodiversity monitoring?

In Madagascar, the protected area (PA) network has undergone substantial expansion in the past two decades in response to rapid forest loss (Gardner et al., 2018). Biodiversity monitoring will be critical to assess the biodiversity status within the PA network and plan for future conservation. However, management of PAs can be hindered by data limitations, including data deficiencies in population dynamics, trends over time, and distributions across large areas for many species of concern (Kremen et al., 2008). Data limitations may be reduced by employing community members to conduct biodiversity surveys, provided that the surveys produce equivalent information to those conducted by scientists. Across the PAs, surveys conducted by CMs recorded mammal species richness that nearly equaled that recorded by scientists. However, for other taxonomic groups, scientists recorded higher species richness than CMs in most PAs. This trend was particularly evident for birds, a highly diverse group. More time spent training CMs could improve the quality of their biodiversity surveys for select taxonomic groups for which differences between CM and scientist surveys were large enough to potentially impact conservation outcomes.

#### 4.1.1. Decision analysis for conservation planning

By quantitatively valuing priorities of decision-makers, our decision analysis provides a method to identify the optimal number of surveys conducted by scientists and CMs and to compare the potential value of different monitoring plans within a given conservation philosophy. Priorities vary across a spectrum of conservation philosophies from an ecocentric philosophy, which prioritizes accurate observation of species, to a people-centered philosophy, which prioritizes incorporating community members in biodiversity monitoring even if at the sacrifice of some accuracy to observations. Despite variation regarding the relative number of species observed by scientists and CMs, decision-analysis showed that integrating community-based monitoring into conservation efforts could provide a financially viable solution that maintains or improves the overall value of monitoring efforts in PAs across the country. This result was consistent across a wide spectrum of conservation philosophies. Even in Beza Mahafaly where scientists recorded on average 23 more bird species than CMs, the ecocentric philosophy required a monitoring plan consisting of 56% CM surveys to attain the optimal value at the lowest cost, while the people-centered philosophy required 97% CM surveys. Optimal monitoring plans often disfavored a large proportion of surveys conducted by scientists because scientists receive larger salaries than CMs and their travel costs must be covered, increasing the financial burden of biodiversity monitoring. It was therefore more cost effective to accumulate species observations over a greater number of CM surveys, only including scientist surveys when CM surveys do not record an equivalent amount of biodiversity. Therefore, if the priority is to monitor common species typically observed during surveys by CMs or a particular species of high conservation concern while including CMs in monitoring efforts, surveys conducted by only community members will suffice. The benefits of such a plan could be great as evident by the high value of a plan consisting of only CM surveys for a people-centered conservation philosophy.

However, if the priority is exhaustive biodiversity monitoring to best estimate species richness, surveys conducted by scientists will be necessary, particularly for birds and reptiles. This scenario can be described as an ecocentric philosophy where high values are only attained when scientists conduct at least some of the surveys. For example, monitoring plans in Beza Mahafaly and Isalo that consist of only CM surveys have lower values than plans containing scientist surveys. Additionally, lower travel costs for scientists, as evident in Ranomafana, may increase the number of scientist surveys recommended to optimize the overall value of monitoring efforts. Ultimately, a decision analysis framework can help identify the optimal number of scientist and CM surveys for any set of monitoring goals and conservation philosophies.

### 4.2. Community involvement in conservation, in Madagascar and around the globe

Community-based monitoring efforts in Madagascar and around the globe support our findings that CM surveys positively contribute to the conservation value of biodiversity monitoring if communities are also involved in resource management and decision-making (Fernandez-Gimenez et al., 2008). In several sites around Madagascar, including the Alaotra Wetlands and Beza Mahafaly Special Reserve, integrating local communities into conservation diversified sustainable income opportunities, increased trust and collaboration with NGOs and governmental agencies, and improved natural resource regulation (Andrianandrasana et al., 2005; Humber, Godley, et al., 2017; Mansourian et al., 2016; Poudyal et al., 2018; Ranaivonasy, Ratsirarson, Mahereza, et al., 2016). In these locations, local community management discouraged illegal use of natural resources through community social pressures, which for Beza Mahafaly has maintained intact forest with robust lemur, bird, and reptile populations (Rahendrimanana et al., 2016; Ranaivonasy, Ratsirarson, Rasamimanana, et al., 2016; Sussman et al., 2012) and for Alaotra led to designation as a Ramsar site and IUCN protected area (Andrianandrasana et al., 2005; Humber, Godley, et al., 2017). For Mitsinjo, a collaborative venture initiated between local community members and foreign scientists in Eastern Madagascar, complete transfer of management rights to the community led to locally organized monitoring efforts that empowered local villages. This impact was exemplified by their ability to obtain funding for conservation activities, conduct knowledge exchange workshops between local and international partners, and establish ecotourism as an additional income source (Dolch et al., 2015). Importantly, successful community-based conservation efforts typically includes natural resource use regulation and enforcement through governmental agencies or local committees (e.g., *dinas* in Madagascar), which establish local legislation and corresponding penalties (Dolch et al., 2015; Ranaivonasy, Ratsirarson, Mahereza, et al., 2016). Consequences of unregulated access to protected areas and forests in Madagascar have included vulnerability of the critically endangered radiated tortoise to the international pet trade (Walker et al., 2014) and threats to forests from small-scale gold mining and logging (Ballet et al., 2019; Cabeza et al., 2019). Decision analysis tools should thus consider financial costs of regulation relative to the savings associated with community led monitoring as well as the possibility for long-term declines in the value of community-based monitoring efforts if regulatory measures are not established.

Community-based monitoring has been shown to provide similar potential for conservation benefits in many locations beyond Madagascar (Conrad & Hilchey, 2011). Synthesizing insights from over 3,000 papers across 126 environmental monitoring schemes globally, Danielsen et al. (2014) found that community-based monitoring can help advance 63% of all Convention on Biological Diversity 2020 indicators. Community-based research also has advanced priorities beyond ecological conservation, such as providing multiple paradigms to understand social and environmental risk factors for public health (Israel et al., 1998). However, successful CBM typically requires establishing monitoring plans in collaboration with local communities for environmental resources that are important to them (Danielsen, Jensen, et al., 2014). Across such diverse contexts, decision analysis provides a useful tool to synthesize stakeholder goals and values and evaluate potential courses of action that consider impacts beyond just the observations community members collect (McGowan et al., 2015).

The mathematical framework we present here is a simplistic representation of the complex costs and values associated with community-based monitoring and does not sufficiently capture the diverse effects of CBM on conservation and local communities. For example, the value of community engagement may increase relative to the number of CM surveys conducted when local communities depend upon the monitored resources or when monitoring efforts are designed to support entire communities rather than individuals (Danielsen et al., 2003; de Araujo Lima Constantino et al., 2012). When developing a framework for decision analysis, complex issues such as these should be discussed with local stakeholders to provide a more complete representation of the attributes that contribute to the overall value of monitoring efforts to conservation and communities. However, the disparate conservation philosophies we consider suggests that integrating community-based monitoring with scientist surveys offers a financially viable method to consistently monitor biodiversity and benefit local communities in the face of limited funding opportunities and global challenges (Razanatsoa et al., 2021). International conservation commitments, including the Aichi Targets presented by the Convention on Biological Diversity, are increasingly calling for the integration of local capacity in management efforts (Convention on Biological Diversity (CBD), 2010). Quantifying the value of alternative monitoring methods will aid decision-makers in developing effective adaptive management plans that can address conservation challenges in dynamic ecological, social, and political landscapes.

## Acknowledgements

We would like to acknowledge the many scientists and local community members who participated in collection of biodiversity data, and continue to maintain and advocate for protected areas in Madagascar.

## Author contributions

DHK, LGS, LR, and BRS conceived the ideas and designed methodology; LR collected the data; LGS analyzed the data; LR and BRS funded the research; LGS led the writing of the manuscript. All authors contributed critically to the drafts and gave final approval for publication.

## Data availability statement

Data are available at the following Zenodo DOI: 10.5281/zenodo.10051226

