## Supplemental Files for "Decision analysis shows scientific and economic value of community-based monitoring in Madagascar"

### Supplementary material

Table S1. The total number of scientist and community member surveys conducted for each taxonomic group and in each park during the years 2019 and 2020.

| Taxon | Scientist or community | Ranomafana | Andringitra | Andohahela | Beza Mahafaly | Isalo | Tsimanampetsotsa |
| --- | --- | --- | --- | --- | --- | --- | --- |
| amphibian | scientist | 42 | 19 | 4 | 2 | 3 | 2 |
| amphibian | community | 27 | 30 | 39 | 33 | 19 | 18 |
| bird | scientist | 47 | 36 | 24 | 45 | 44 | 40 |
| bird | community | 179 | 142 | 209 | 173 | 105 | 169 |
| mammal | scientist | 28 | 20 | 8 | 35 | 18 | 2 |
| mammal | community | 193 | 83 | 100 | 168 | 147 | 144 |
| reptile | scientist | 33 | 28 | 19 | 45 | 38 | 40 |
| reptile | community | 170 | 125 | 213 | 172 | 163 | 161 |

Table S2. Cost of salary and travel for community and scientist surveys. For each park, cost per survey was calculated as the total amount spent on salaries divided by the number of surveys that were conducted by scientists or community members.

|  | Ranomafana | Andringitra | Andohahela | Beza Mahafaly | Isalo | Tsimanampetsotsa |
| --- | --- | --- | --- | --- | --- | --- |
| Scientist |  |  |  |  |  |  |
| *cost/survey (USD)* | 16.15 | 19.00 | 23.75 | 19.95 | 17.10 | 19.00 |
| *Travel cost (USD)* | 57.42 | 155.28 | 938.07 | 105.85 | 86.58 | 214.91 |
| Community member |  |  |  |  |  |  |
| *cost/survey (USD)* | 0.31 | 0.48 | 0.58 | 0.45 | 0.44 | 0.57 |
| *Travel cost (USD)* | 0 | 0 | 0 | 0 | 0 | 0 |

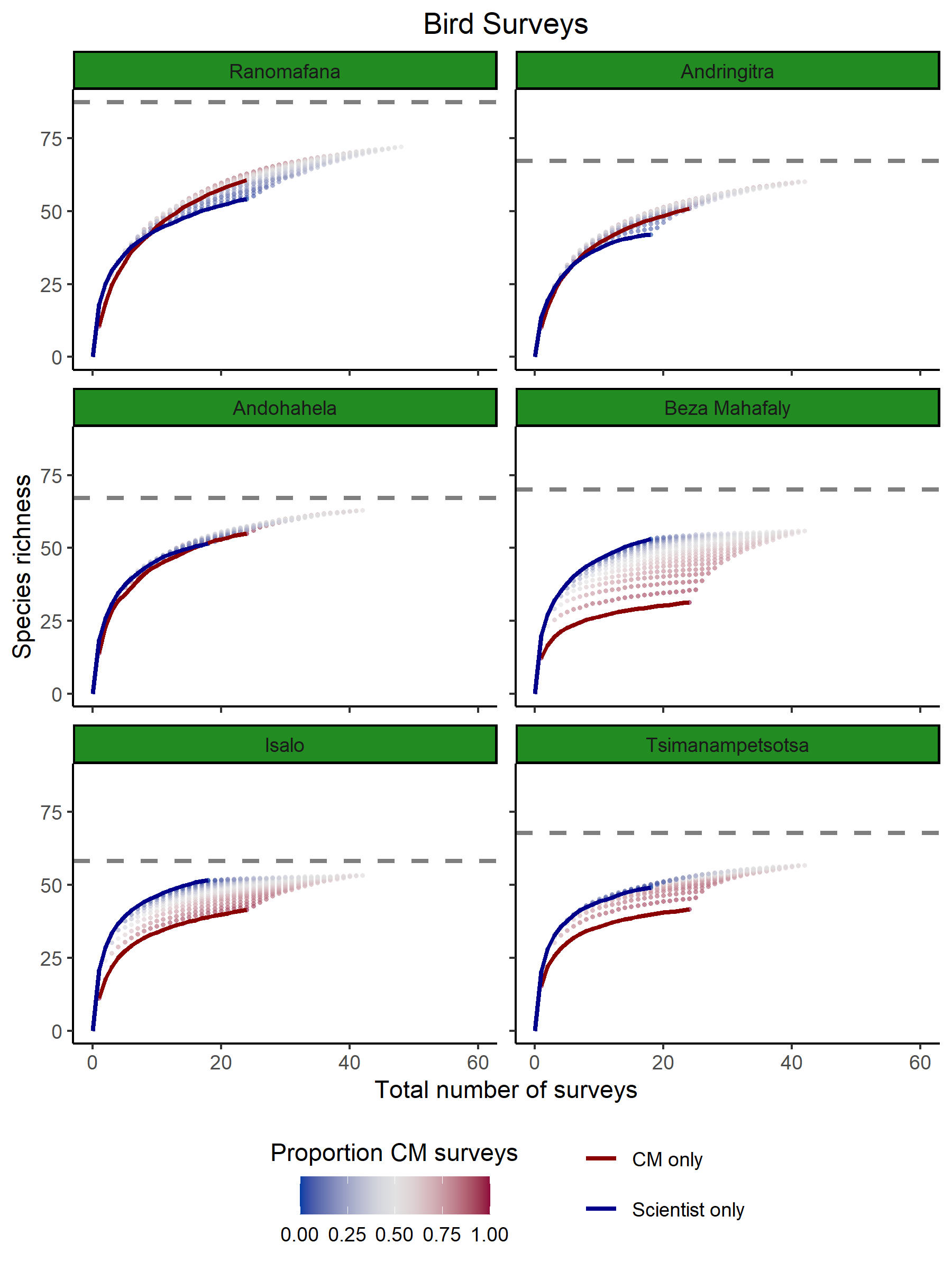

Fig. S1. Species accumulation for birds in six PAs across the total number of surveys conducted by a combination of scientists and community members (CMs). Each point represents a given number of scientist and CM surveys and is colored by the proportion of CM surveys out of the total number of surveys conducted. surveys of vertebrates in six protected areas. The blue line represents species accumulation for scientists only and the red line represents species accumulation for CMs only. The gray dashed line represents the estimated total number of species in each PA.

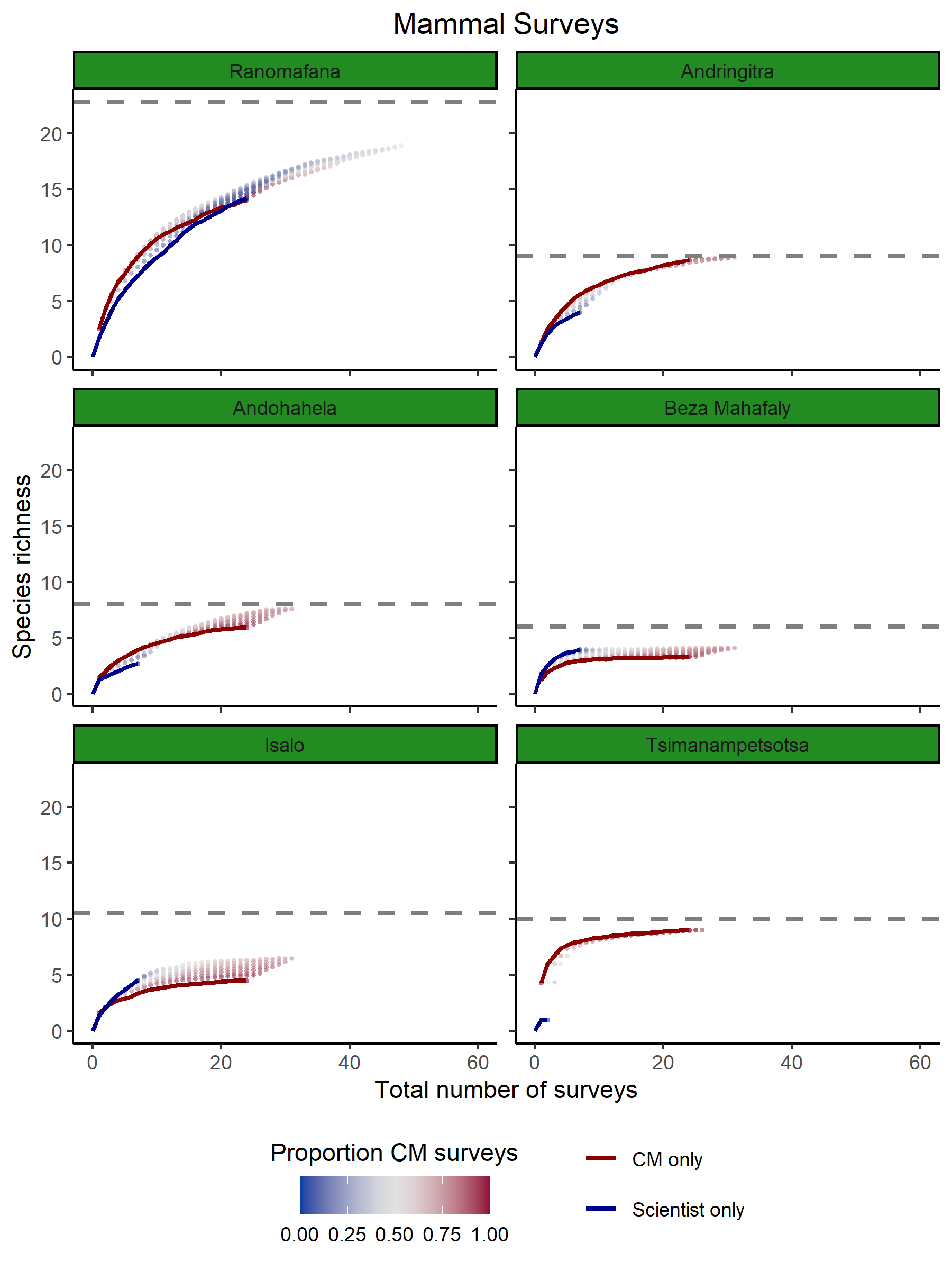

Fig. S2. Species accumulation for mammals in six PAs across the total number of surveys conducted by a combination of scientists and community members (CMs). Each point represents a given number of scientist and CM surveys and is colored by the proportion of CM surveys out of the total number of surveys conducted. surveys of vertebrates in six protected areas. The blue line represents species accumulation for scientists only and the red line represents species accumulation for CMs only. The gray dashed line represents the estimated total number of species in each PA.

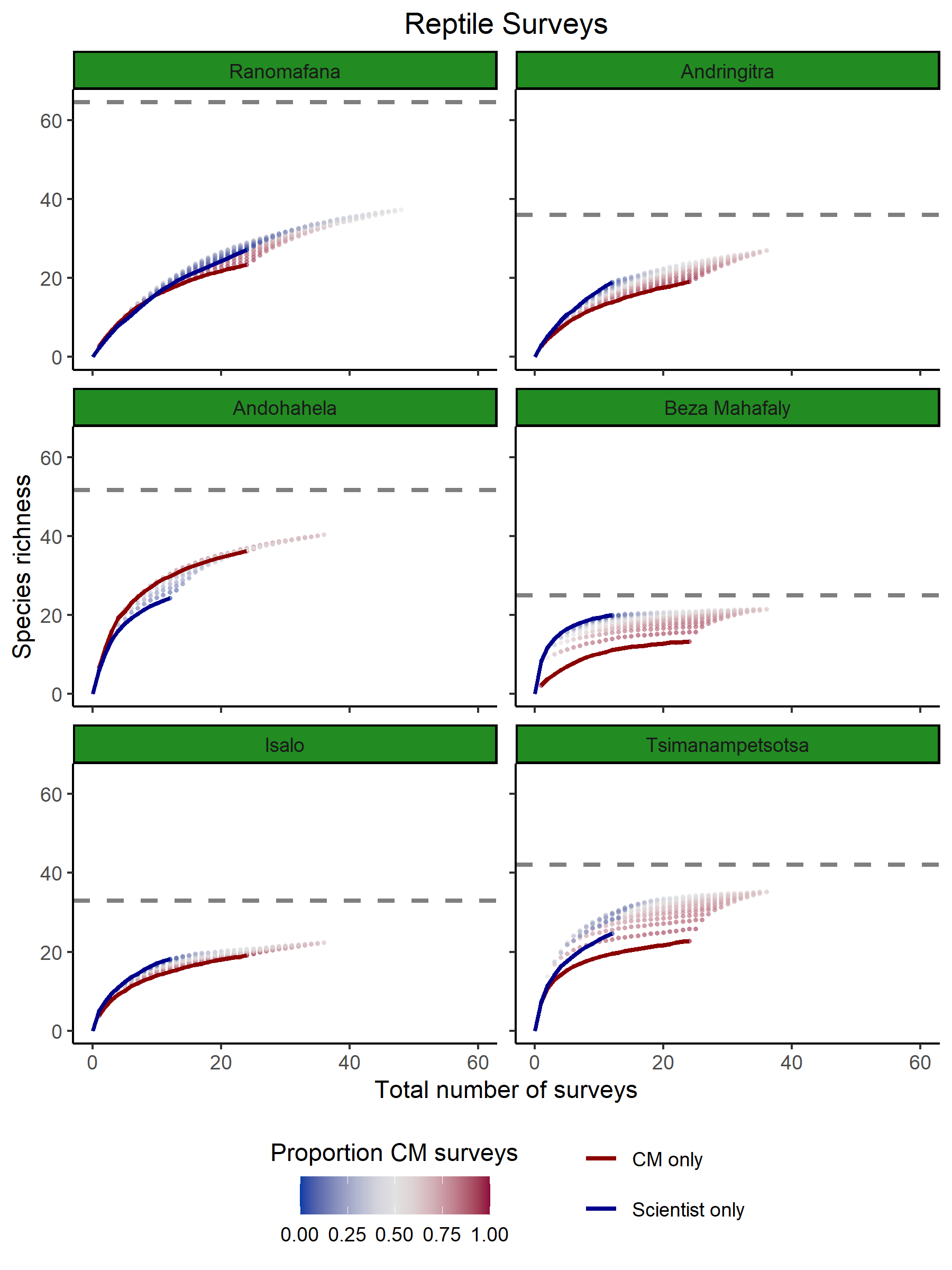

Fig. S3 Species accumulation for reptiles in six PAs across the total number of surveys conducted by a combination of scientists and community members (CMs). Each point represents a given number of scientist and CM surveys and is colored by the proportion of CM surveys out of the total number of surveys conducted. surveys of vertebrates in six protected areas. The blue line represents species accumulation for scientists only and the red line represents species accumulation for CMs only. The gray dashed line represents the estimated total number of species in each PA.

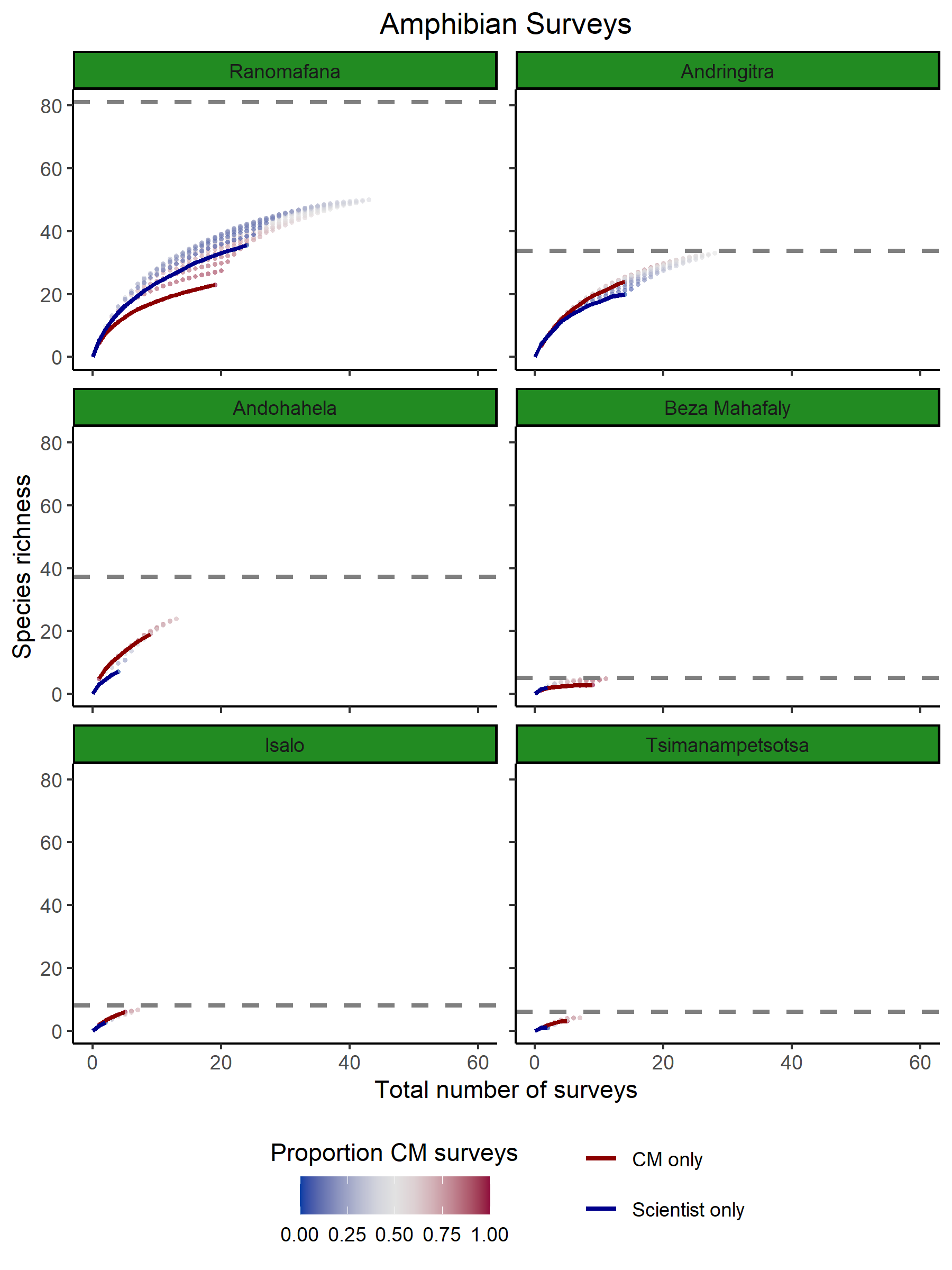

Fig. S4. Species accumulation for amphibians in six PAs across the total number of surveys conducted by a combination of scientists and community members (CMs). Each point represents a given number of scientist and CM surveys and is colored by the proportion of CM surveys out of the total number of surveys conducted. surveys of vertebrates in six protected areas. The blue line represents species accumulation for scientists only and the red line represents species accumulation for CMs only. The gray dashed line represents the estimated total number of species in each PA.

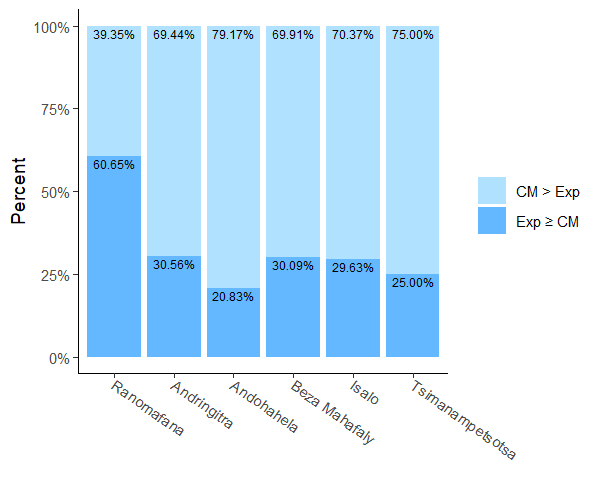

Fig. S5. The percentage of conservation philosophies in which the ideal number of CM and scientist surveys needed to obtain the optimal value at minimum financial cost consisted of more CM surveys (lighter blue) or more scientist surveys (darker blue). The optimal value refers to the maximum possible value minus 0.01, which represents that amount overall value decision-makers are willing to compromise to reduce the financial cost of monitoring efforts.

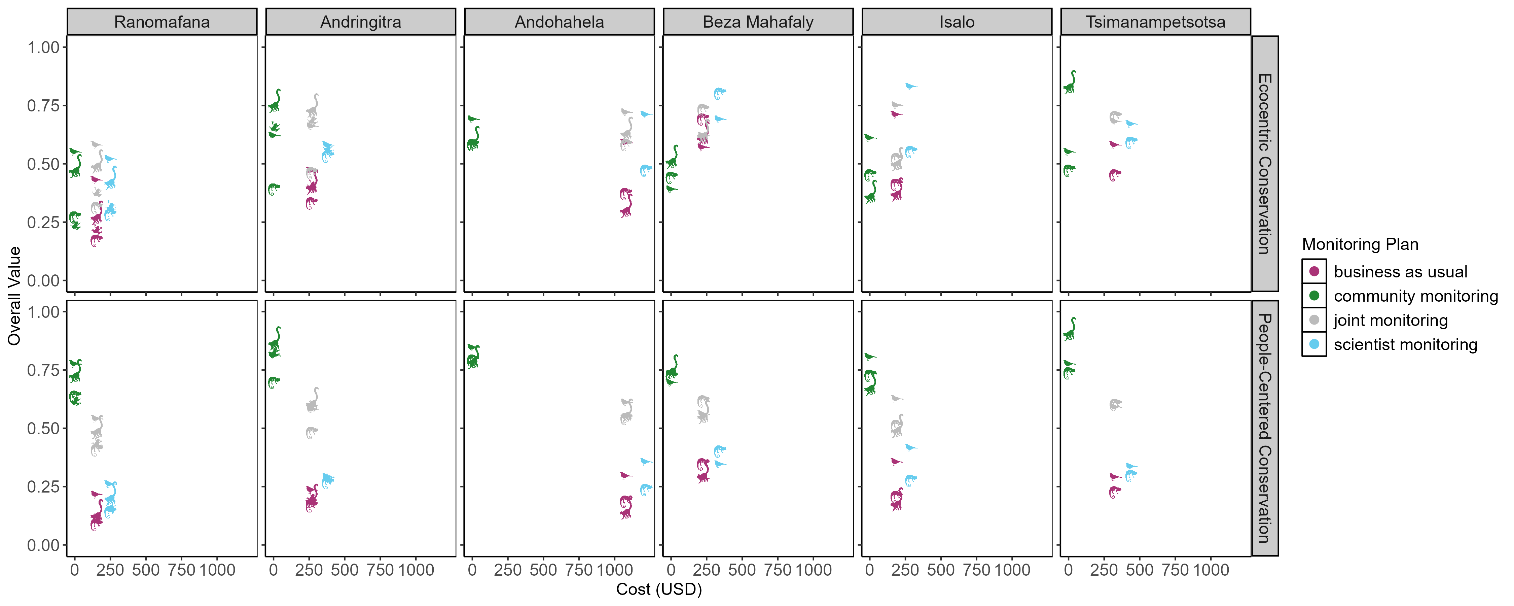

**Fig. S6.** Overall costs and values associated with four monitoring plans for two different conservation philosophies. The ecocentric conservation philosophy is defined by a species richness weight of 1, a linear species richness function, and a community engagement weight of 0. The people-centered conservation philosophy is defined by a species richness weight of 0.5, a linear species richness function, a community engagement weight of 0.5, and a linear community engagement function. The four monitoring plans include business as usual (6 scientist surveys), community engagement (6 scientist and 6 CM surveys), community monitoring (12 CM surveys), and scientist monitoring (12 scientist surveys). Overall monitoring costs and values are calculated per PA and taxon (represented by point shape – amphibian (frog), bird, mammal (lemur), and reptile (chameleon)). If the monitoring plan contains more surveys that were conducted during the study (six or twelve surveys depending on the monitoring plan), then it was excluded from the analysis.
